# Evidence associating neutrophilia, lung damage, hyperlactatemia, blood acidosis, impaired oxygen transport, and mortality in critically ill COVID-19 patients

**DOI:** 10.1101/2023.09.17.558185

**Authors:** Basma A. Yasseen, Aya A. Elkhodiry, Hajar El-sayed, Mona Zidan, Azza G. Kamel, Rehab Hamdy, Hend E. El-Shqnqery, Omar Samir, Ahmed A. Sayed, Mohamed A. Badawy, Aya Saber, Marwa Hamza, Riem M. El-Messiery, Mohamed El Ansary, Engy A. Abdel-Rahman, Sameh S. Ali

**Author notes:** Equal contributions (Basma A. Yasseen, Aya A. Elkhodiry). Corresponding authors: Prof. Sameh S. Ali; or Dr. Engy A. Abdel-Rahman. Research Department, Children’s Cancer Hospital Egypt 57357, Cairo, Egypt, Ext-7233.

## Abstract

**Background:** COVID-19 severity and high in-hospital mortality are often associated with severe hypoxemia, hyperlactatemia, and acidosis. Since neutrophil numbers in severe COVID-19 can exceed 80% of the total circulating leukocytes and that they are massively recruited to infected lungs, we investigated whether metabolic acidosis mediated by the glycolytic neutrophils is associated with lung damage and impaired oxygen delivery in critically ill patients.

**Methods:** Based on prospective mortality outcome, 102 critically ill-hospitalized COVID-19 patients were divided into two groups: ICU-Survivors (ICU-S, n=36) and ICU-Non-survivors (ICU-NS, n=66). Blood samples were collected from patients and control subjects to explore correlations between neutrophil counts, lung damage, glycolysis, blood lactate, blood pH, hemoglobin oxygen saturation, and mortality outcome. We also interrogated isolated neutrophils for glycolytic activities and for apoptosis using high-throughput fluorescence imaging complemented with transcriptomic analyses. Stratified survival analyses were conducted to estimate mortality risk associated with higher lactate among predefined subgroups.

**Results:** Neutrophil counts were consistently higher in critically ill patients while exhibiting remarkably lower apoptosis. Transcriptomic analysis revealed miRNAs associated with downregulation of genes involved in neutrophils apoptosis. Both CT lung damage scores and neutrophil counts predicted mortality. Severinghaus fitting of hemoglobin oxygen saturation curve revealed a right-shift indicating lower oxygen capacity in non-survivors, which is consistent with lower blood-pH observed in the same group. Levels of blood lactate were increased in patients but significantly more in the ICU-NS relative to the control group. ROC analysis followed by Kaplan-Meyer survival analysis stratified to the obtained cut-off values showed that CT damage scores, neutrophil counts, and lactate levels are predictors of mortality within 15 days following blood collection.

**Conclusion:** The current results implicate neutrophilia as a potential player in metabolic acidosis and deranged oxygen delivery associating SARS-CoV-2 infection thus contributing to mortality outcome.

## Introduction

The world-meter of confirmed cases and mortality counts related to the coronavirus disease 2019 (COVID-19) pandemic has been ticking like an eternal clock since December 2019! While writing this report the world was once again alerted to the highly mutated variant BA.2.86, whose mutations could amount to an evolutionary jump matching the emergence of the Omicron variant in 2021. Earlier this year, COVID surged in China with the highly contagious XBB.1.5 variant, which was declared as a variant of ‘growth advantage’ by the World Health Organization indicating that the battle with the pandemic is not over. Lack of effective treatments, ineffective global vaccine coverage, and rapid evolution of viral variants and pathogenicity are all daunting characteristics of the battle with SARS-CoV-2 virus. Despite increasingly inhabited areas of research on COVID-19 pathology, much wider territories remain uncharted and await further explorations. One outstanding question is how disproportionately upregulated and downregulated immune cells interact with the lung and orchestrate the inflammatory response during the course of infection particularly in critically ill patients.

Neutrophilia, lung damage, hypoxia and lactic acidosis are main defining hallmarks of severe COVID-19 pathology. Neutrophilia, or the remarkable increase in the number of neutrophils in the bloodstream, is reported in up to 70% of COVID-19 patients and is associated with more severe symptoms [1]. The pathology of COVID-19 is also associated with systemic priming and delayed apoptosis/clearance of neutrophils from the lungs [2]. Neutrophils activation is a double-edged sword during the immune response to infections as they act as effector cells that are fundamental components of innate immunity but are also profoundly histotoxic [3,4]. Neutrophils are quickly and massively recruited by the pulmonary system to accumulate in high abundance in the lung interstitium and alveolar airspace of patients with COVID-19 [5]. It has been proposed that activated populations of immune cells including neutrophils and macrophages mobilized into the lungs, constitute a major source of lactate released by the lung during septic shock [reviewed in [6]], especially when sepsis is complicated with acute respiratory distress syndrome (ARDS) [7].

Hypoxemia is also common in COVID-19 patients and is associated with increased mortality [8]. Because tissue hypoxia provokes a switch to anaerobic glycolysis and enhances lactate production, hyperlactatemia is often considered as an indication of hypoxia in critically ill patients. Hypoxia and acidosis reinforce each other in a viscous cycle; a drop in blood plasma pH by less than 0.2 units was found to cause a significant reduction in the ability of hemoglobin to bind oxygen [9]. Conversely, lactic acidosis, or the increase in the amount of lactic acid in the bloodstream, is associated with increased morbidity and mortality [10]. Indeed, it has been demonstrated that blood lactate values are higher in patients with a severe form of COVID-19 than in patients with milder illness [11–13], and amongst hospitalized patients, lactate values were found to be the highest in non-survivors [reviewed in: [14]]. However, Iepsen et al. argued that lactate is unrelated to tissue hypoxia but is likely to reflect mitochondrial dysfunction and high adrenergic stimulation [15]. Studies exploring metabolic remodeling of immune cells in the context of severe COVID-19 are thus needed to settle this debate and to allow better care in the ICU for patients with COVID-19, ARDS, and sepsis.

In a recent study, we demonstrated that neutrophil-mediated oxidative damage of serum albumin is an accurate predictor of COVID-19 mortality [16]. Since albumin is also highly sensitive to pH changes, we asked if metabolic rewiring to a more lactate-producing glycolytic phenotype in leukocytes in general and neutrophils in particular contributes to the observed protein structural modification. Lactate is a main byproduct of glycolysis fueled by human neutrophils [17]. Neutrophils are classically viewed to rely on glycolytic metabolism for energy production under basal conditions, where pyruvate is transformed into lactate for regenerating NAD+ during glycolysis. On the other hand, mitochondria in neutrophils have been usually viewed as minor bioenergetic sources and their role has been restricted to the initiation of apoptosis [18]. In studies addressing metabolic rewiring in severe COVID-19 patients, conflicting observations regarding neutrophils metabolism have been reported. For example, in severe patients, neutrophils were found to display enhanced glycolysis and defective mitochondria function [19]. Moreover, McElvaney et al. [20] have shown elevated cytosolic levels of an enzyme (pyruvate kinase M2) that catalyzes the last glycolytic step, as well as lactate in neutrophils isolated from severe COVID-19 patients, hinting at enhanced glycolytic flux. On the other hand, a study performed on neutrophils isolated from COVID-19 patients with ARDS demonstrated a diminished glycolytic flux and a shift to oxidative metabolism [21]. The present study was originally planned to explore neutrophils’ metabolic profile in relation to mortality outcome in a cohort of severe COVID-19 patients and was expanded to explore the contribution of neutrophils to lung damage, blood acidosis, and reduced oxygen carrying capacity of hemoglobin in those patients.

## Methods

### Demographic, clinical, and hematologic characteristics of the studied COVID-19 cohort

This is a prospective observational cohort study conducted to assess patients who tested positive for COVID-19 through RT-PCR test. Patients recruitment took place at the ICU facility within Kasr Alainy Cairo University Hospital-Internal Medicine Quarantine Hospital. Standard supportive therapy, including supplemental oxygen and symptomatic management, was administered as clinically indicated. Patients exhibiting moderate to severe hypoxia, defined as requiring a fraction of inspired oxygen (FiO2) of ≥40%, were transferred to the intensive care unit (ICU) for escalated care, including invasive mechanical ventilation when deemed necessary. Within the ICU patient cohort, individuals were categorized into two groups based on their subsequent mortality outcomes: those who survived (ICU-S) and those who died within 20 days of sample collection (ICU-NS). All available samples and analysis were conducted by blinded operators and were included in the final dataset. In adherence to the ethical principles outlined in the Declaration of Helsinki, written informed consents were obtained from all study participants. The study’s design and protocol underwent thorough evaluation and received approval from the Institutional Review Board (IRB) at the Children’s Cancer Hospital. The study was assigned IRB number 31-2020, initially issued on July 6, 2020, and subsequently renewed on July 28, 2021.

The demographic and clinical characteristics of the studied patients are given in Table 1. Participants were divided into two groups: ICU-Survivors (ICU-S, n=36) and ICU-Non-Survivors (ICU-NS, n=66). The two groups did not exhibit statistically significant differences between most of the clinical or demographic characteristics except for the blood saturation level (p=0.045). Additionally, the number of patients suffering from diabetes is significantly higher in the ICU-NS group (p = 0.008). While, the number of patients who were treated with steroids, remdesivir, hydroxychloroquine and/ or carbapenem antibiotics was insignificantly different in both groups.

**Table 1:** Demographic and clinical characteristics of the participants.

|  | ICU-S n (%) | ICU-NS n (%) | p value |
| --- | --- | --- | --- |
| Male | 23 (63.9%) | 34 (51.5%) | 0.23 |
| Age | 61 (25-77) | 66 (17-98) | 0.12 |
| sO<sub>2</sub> | 94 (59-99) | 90 (41-97) | **0.045** |
| Diabetes | 6 (16.7%) | 28 (42.4%) | **0.008** |
| Cardiovascular diseases | 11 (30.6%) | 26 (39.4%) | 0.37 |
| Cancer | 4 (11.1%) | 3 (4.5%) | 0.21 |
| Asthma | 2 (5.6%) | 3 (4.5%) | 0.82 |
| Insulin | 6 (16.7%) | 16 (24.2%) | 0.37 |
| Anticoagulant | 16 (44.4%) | 34 (51.5%) | 0.49 |
| Steroids | 11 (30.6%) | 31 (47%) | 0.11 |
| Hydroxychloroquine | 1 (2.87%) | 2 (3%) | 0.94 |
| IL6 inhibitors | 3 (8.3) | 5 (7.6%) | 0.89 |
| Remdesivir | 3 (8.3%) | 15 (22.7%) | 0.07 |
| Ivermectin | 1 (2.8%) | 2 (3%) | 0.94 |
| carbapenem antibiotics | 11 (30.6%) | 32 (48.5%) | 0.08 |
| Fluoroquinolone | 7 (19.4%) | 21 (31.8%) | 0.18 |
| Oxazolidinone antibiotic | 7 (19.4%) | 10 (15.2%) | 0.58 |

Table 2 shows the laboratory results of the participants. When comparing all parameters in the two groups, we observed changes following the same reported trends in our previous studies [16,22]. Non-significant changes were reported in all of the parameters except for a significant decrease in albumin level in the ICU-NS group (p =0.03) and in platelets count (p= 0.017). The results also show a significant increase in the white blood cells count (p =0.02), C-reactive protein (p=0.008) and D-Dimer level (p=0.008) in non-survivors.

**Table 2:** Laboratory results of participants.

|  | ICU-S | ICU-NS | P-value |
| --- | --- | --- | --- |
| WBCs ($\times 10^3$ /ml) | 9.5 (4.4-20.7) | 13.2 (4.2-47) | **0.02** |
| Platelets ($10^6$ /ml) | 274.5±115 | 211.26±118 | **0.017** |
| Lymphocytes | 3.45 (0.48-24.5) | 3.4 (0.25-32.5) | 0.92 |
| Monocytes | 0.99 (0.11-10.2) | 1.26 (0.2-12.4) | 0.44 |
| INR | 1.12 (1-2.86) | 1.2 (1-4.4) | 0.34 |
| CRP (mg/L) | 56.5 (1.9-265) | 109.03 (12-305) | **0.008** |
| D-Dimer (mg/ml) | 0.98 (0.17-5.2) | 2 (0.19-19.1) | **0.008** |
| IL-6 (pg/ml) | 25 (2.3-1830) | 32.4 (1.77-4500) | 0.90 |
| Ferritin | 797 (218-2032) | 1148 (139-20000) | 0.06 |
| Albumin | 2.75 (2.1-3.8) | 2.5 (1.6-246) | **0.03** |
| haemoglobin (g/dl) | 11.7 (5.7-15.2) | 11.2 (5.5-74) | 0.68 |
| ALT (U/L) | 19 (8-92) | 35 (3-232) | 0.23 |
| AST (U/L) | 29 (17-40) | 37 (12-168) | 0.08 |
| Creatinine (mg/dl) | 0.96 (0.3-6.85) | 1.3 (0.38-6.9) | 0.13 |

### Radiologic assessment of lung damage

A lung CT damage score for each patient was calculated according to a semi-quantitative CT severity scoring protocol proposed by Pan et al. [23]. This scoring system is based on the extent of parenchymal involvement per each of the 5 lobes [24] as follows: (0) no involvement; (1) <5% involvement; (2) 5-25% involvement; (3) 26-50% involvement; (4) 51-75% involvement; or (5) >75% involvement. The resultant total CT score is the sum of the obtained lobar scores and ranges from 0 to 25.

### Blood sample collections, handling, and processing for blood cells isolation and analyses

A volume of 10 mL of fresh blood was collected from all subjects on Acid Citrate Dextrose (ACD) tubes (Greiner Bio-One GmbH, Kremsmünster, Austria). For flow cytometry measurements, 2 mL of collected citrated whole blood was incubated for 15 min with RBCs lysis buffer composed of NH_4_Cl (ammonium chloride), NaHCO_3_ (sodium bicarbonate), and EDTA (disodium) [25]. The rest of the blood -(8 mL) was used for blood cells isolation as previously described [26]. Briefly, whole blood was centrifuged (300×g) at 25 °C for 15 min with no brakes. (1) The upper clear layer (platelet rich plasma) was transferred to a new tube and centrifuged to get platelets pellet and platelets poor plasma as described previously [22]. (2) The lower layer was gently layered over equal volumes of lymphocyte separation medium (1.119 and 1.077). Tubes were then centrifuged at 500×g for 35 min at 25 °C with no brakes and lowered acceleration. After centrifugation, the second layer containing peripheral blood mononuclear cells (PBMCs) was collected and washed in PBS, and the pellet was suspended in 100 µl PBS. The layer comprising neutrophils was collected and washed in Hank’s Balanced Salt Solution (HBSS), then centrifuged again at 350×g for 10 min at 25 °C. The supernatant was discarded, the pellet was suspended in RBCs lysis buffer and incubated at room temperature for 15 min. Following centrifugation for 10 min at 350×g, the pellet was suspended in 100 µl PBS. Neutrophils and PBMCs counts were performed manually by a hemocytometer. Cellular viability was monitored for each sample using trypan blue, and only cells exhibited more than 90% viability were used in the analysis.

### Assessment of neutrophil counts and neutrophils apoptosis by flow cytometry

Suspended lysed blood cells were centrifuged at 500xg for 5 minutes and washed twice with phosphate-buffered saline (PBS). Cells were then suspended in PBS, and the detection of neutrophils was carried out using 13-color flow cytometer CytoFLEX system (Beckman Coulter Life Sciences CytoFLEX benchtop flow cytometer) as follows. Suspended cells were incubated in the dark for 30 min at room temperature with CD66b-APC-Alexa Fluor 750 (Beckman Coulter Life Sciences, B08756) for the detection of neutrophils. Cells were washed with PBS by the end of the incubation period and then suspended in 300 μl PBS and examined by flow cytometry for gating neutrophil-specific CD66b-APC-Alexa Fluor 750 positive population. For evaluation of neutrophil cell death, the cells were washed with PBS and resuspended in 50 µl of 1X annexin binding buffer containing 2 µl of FITC-annexin V (Miltenyi Biotec, 130-092-052). Cells were then incubated at room temperature for 30 min in the dark. After incubation, the cells were washed with PBS and suspended in 300 µl of 1X annexin binding buffer. A total of 20,000 events were acquired and then analysed using CytExpert software.

### Measurement of apoptosis in isolated neutrophils by fluorescence imaging

Neutrophils cell death was detected and quantified using the annexin V dye (Thermo Fisher Scientific, A23202). Isolated cells were seeded and incubated for 30 min at 37°C followed by centrifugation to form a monolayer. Cells were then stained with Annexin V in 1x annexin binding buffer as per the manufacturer’s recommendations for 30 min. Images were acquired (1-5 images per subject) using the Cytation 5 Cell Imaging MultiMode Reader (Agilent). Images were processed to quantify fluorescence intensity using Gen5 software package 3.08.

### Measurement of Extracellular Acidification Rate (ECAR) in different populations by Seahorse XF96 Flux Analyzer

Extracellular acidification rates (ECAR) in neutrophils, PBMCs, and platelets were assessed using XF analysis (XF96, Seahorse analyzer, Agilent) as described previously [22]. Cells were plated on 96 well format XF plates at a density of 2 × 10^5^ neutrophils/well, 2.5 × 10^5^ PBMCs/well and 20 × 10^6^ Plts/well in unbuffered DMEM deprived of glucose and pyruvate (DMEM; with 4 mM L-glutamine, pH 7.4 at 37°C) for the glycolytic stress tests. Plates were centrifuged (800xg for 5 minutes) to form cells monolayer in wells. Plates were then incubated for 30-40 min at 37 °C in non-CO2 incubator. A baseline measurement for ECAR was acquired for 30 min. ECAR values were examined following consecutive injections of glucose (5.5 mM), oligomycin (2.5 µM) and 2-deoxy-D-glucose (2-DG) (50mM). Glycolysis, glycolytic capacity, and glycolytic reserve were calculated by subtracting the average rates before and after the addition of glucose, ATP synthase inhibitor oligomycin and 2-DG. All measurements were normalized to seeding density before being multiplied by the respective neutrophils count per µl blood determined by flow cytometry.

### Measurement of L-lactate in plasma

L-lactate level was assessed in plasma using colorimetric assays according to the manufacturer’s protocol with slight modifications (Lactate Assay Kit, Sigma-Aldrich, Missouri, USA, # MAK064-1KT). 25 μl of lactate standards or plasma samples were mixed with an equal volume of Master Reaction Mix (consists of: 92% lactate assay buffer; 4% lactate probe and 4% lactate enzyme mix). Following incubation for 30 minutes, absorbance was measured at 570 nm (A570), using the Cytation 5 Cell Imaging Reader (Agilent BioTek, CA, USA). Lactate levels present in the samples were estimated from the standard curve.

### Measurement of L-lactate in blood

L-lactate level was assessed in the blood using blood lactate meter according to the manufacturer’s protocol (Lactate Pro 2, ARKRAY Inc., Kyoto, Japan).

### Small RNA library preparation and sequencing

RNA libraries were generated using the NEXTFLEX® Small RNA-Seq Kit (PerkinElmer, USA; Cat. No.: NOVA-5132-05). For each library, 400 ng of purified RNA was used as an input for library preparation according to the manufacturer’s instructions. Ligated libraries were reverse transcribed and amplified with a unique barcode primer for each one. DNA fragments ∼ 150 bp (miRNA sequences plus 3′ and 5′ adaptors) were determined using 6% TBE-PAGE gel and then retrieved in a 300μl elution buffer for purification. The size distribution of the pooled library was checked using the Bioanalyzer DNA assay (Agilent, USA; Cat. No.: 5067-1504) and the concentration was assessed using the Qubit dsDNA HS Assay (ThermoFisher Scientific, USA; Cat. No.: Q33230). The final pooled library was sequenced for around 2,400 miRNAs using the Illumina MiSeq system (Illumina, Inc., USA).

### Bioinformatics analysis

Raw data quality was inspected using FastQC [27]. In addition, low-quality reads and adaptors were trimmed using Cutadapt as instructed by the manufacturer [28]. Filtered reads were mapped using Bowtie 1 against the human genome reference GRCh38 (accession number GCA_000001405.29). Mapped reads were quantified using FeatureCounts based on the human miRNA coordinate file (gff) retrieved from the miRbase database release 22.1 [29,30]. Count data were filtered and normalized, and differential expression analysis between survivor and non-survivor groups was performed using DESeq2 package in R V4.1.2 [31]. Differentially expressed miRNAs (DEMs) were selected to have a log2 fold change ˃ 1.5 and an adjusted p-value (Adj. p) value <0.05. DEMs log2 fold change and adjusted p-value were visualized by lollipop plot using the ggplot R package [32]. In addition to heatmap plot for DEMs normalized count using the pheatmap R package [33].

### Identification of miRNA putative gene targets and network analysis

The miRPathDB database was used to perform pathway enrichment analysis for differentially expressed miRNAs against the KEGG database, and only pathways with experimental evidence were selected [34,35]. Enriched pathways were retrieved and the ggplot R package was used to draw a bubble plot representing the number of regulated genes and the p-value retrieved from miRPathDB. Finally, pathway visualization was performed using the Pathview R package representing regulated genes [36].

### Statistical analysis and data presentation

Statistical analysis and data graphing were performed using OriginPro 2022b (OriginLab Corporation, Northampton, USA). Graphical representations of data utilized box-and-whisker plots while showing the actual scatter of the data points combined with a distribution curve to indicate normality. Violin plots were also utilized to outline statistical analyses outcomes and to demonstrate data scatter, distribution, and mean ± SD or SEM, as stated in the figure legends. Density Color Mapping, which assigns color based on the density of points in a two-dimensional scatter plot, is used to infer associative variance and covariance directionality between the two-plotted parameters. Exact p-values are given for each comparison on most graphs and in the text. Following tests for normality (Shapiro–Wilk test), variables that fulfilled the normality test were analyzed using the ANOVA test followed by Tukey post hoc tests to compare the differences between the three groups or the Independent Samples t Test while non-normally distributed data was analyzed using the Mann-Whitney test to compare the differences between the two groups. Categorical variables are reported as counts and percentages while continuous variables are expressed as mean ± SD or a median (range). Differences between percentages were assessed by Pearson’s X2 tests or Fisher exact tests when the number of observations per group was less than 5. The χ2 tests provided results that tested the hypothesis that mortality and a given variable (e.g., sex or comorbidity) are independent. Linear regression analysis and Pearson correlation coefficients were obtained without asymptotic assumptions. When *p* is less than the significant level of 0.05, there is significant evidence of association between mortality and the variable.

## Results

### Remarkable increase in counts and reduction of apoptotic surface markers on neutrophils of severe patients

We started by following changes pertaining to neutrophil counts and apoptotic profiles in relation to mortality outcome in the studied cohort of subjects. First, we used flow cytometry to assess the percentages of neutrophils in all patients (Fig. 1A, B). Similar to our previously published work [16], a significant rise in neutrophil counts (% total) was observable when going from Control (36.1%±13.1%; n=16) to ICU-S (64.9%±19.4%; n=32) to ICU-NS (74.6%±13.4%; n=53) groups (ICU-S vs. Control: p=3.6×10^-8^; ICU-NS vs. Control: p=0; ICU-S vs. ICU-S: p=0.018). These results support the conclusion of a meta-analysis reporting the validity of neutrophilia as a mortality predictor in severe COVID-19 patients [37]. Furthermore, and to shed light on the mechanism of neutrophil accumulation, we evaluated apoptosis in freshly isolated neutrophils from all groups by measuring the percentage of annexin-V^+^ cells using both flow cytometry (Fig. 1C-E) and fluorescence imaging (Fig. 1F, G). Our flow cytometry analyses showed that neutrophils of both COVID-19 groups exhibit significantly diminished annexin-V^+^ populations (early apoptosis) relative to the control group (Control: 45.9% ± 27.4%; n=22, ICU-S: 24.4% ± 16.2%; n=13, ICU-NS: 18.0% ± 17.1%; n=26. ICU-S vs. Control: p=0.01; ICU-NS vs. Control: p= 9×10^-5^; ICU-S vs. ICU-NS: p=0.66). These results were confirmed through the quantifications of median fluorescence intensities (MFI) of individual neutrophil fluorescence images (1-5 images per subject) from controls (n=7), ICU-S (n=6) and ICU-NS (n=11) stained with annexin-V dye (Fig. 1F, G). These data suggest that suppression of neutrophil apoptosis/clearance maybe implicated in the neutrophilia observed in critically ill COVID-19 patients (Control: 6039 ± 2351; n=7, ICU-S: 3421 ± 1773; n=6, ICU-NS: 2782 ± 1440; n=11. ICU-S vs. Control: p=0.04; ICU-NS vs. Control: p= 0.004; ICU-S vs. ICU-NS: p=0.77).

**Figure 1.**
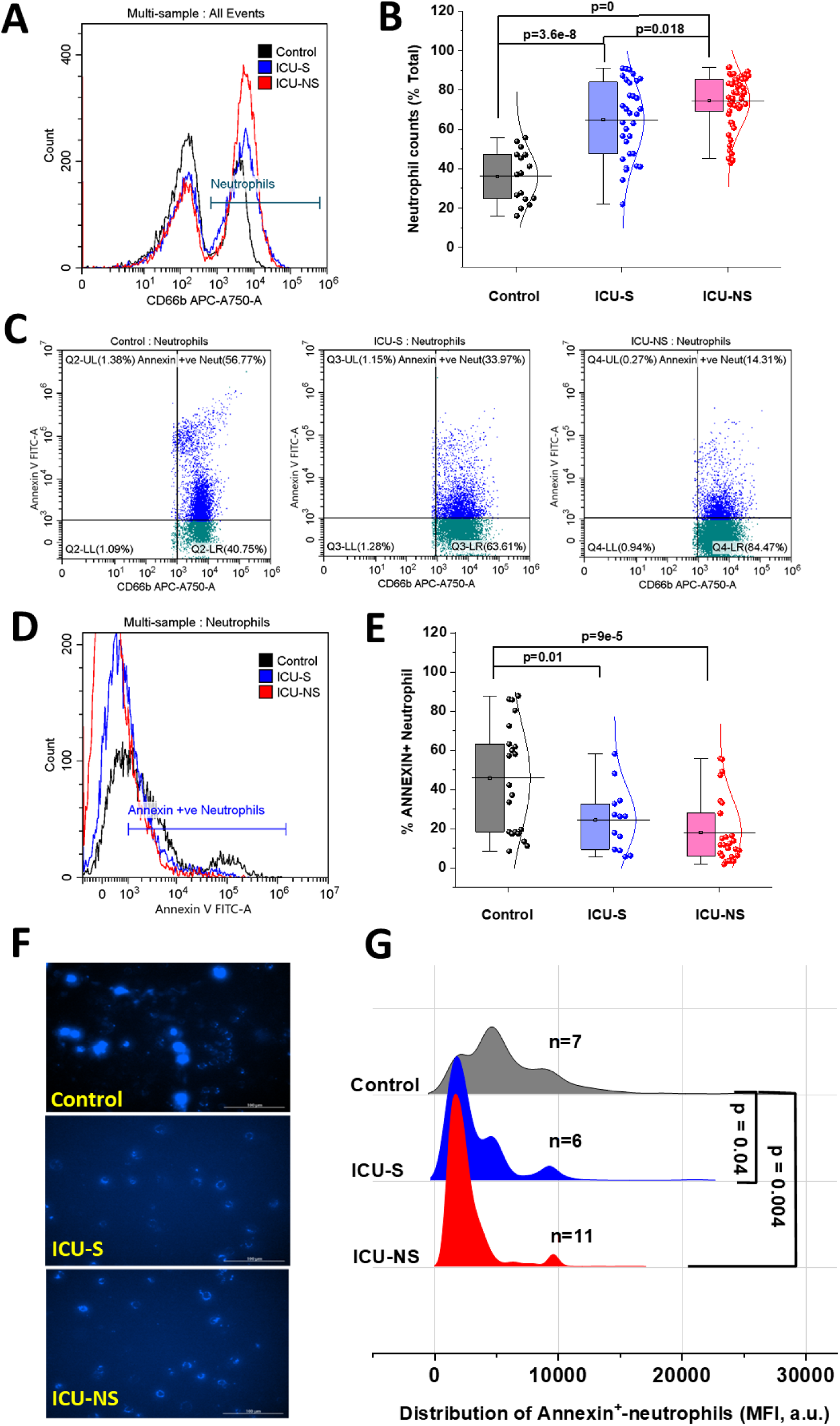
Increased neutrophil counts and decreased apoptotic neutrophils in critically ill COVID-19 patients. A) Representative flow cytometric histograms comparing neutrophil counts in the control, ICU-S, and ICU-NS roups. (B) When neutrophil counts were compared for all groups, both ICU-S and ICU-NS groups displayed tatistically significant increase in neutrophil counts relative to control groups (n=16). Neutrophil counts were lso significantly higher in ICU-NS when compared to ICU-S (n=32, and 53 for ICU-S, and ICU-NS; respectively). C) Representative flow cytometry scatter diagrams of annexin V+ neutrophil populations for all groups. (D) epresentative flow cytometric histograms comparing annexin V+ neutrophils for all groups. (E) Diminished vels of annexin V+ cells in ICU-S and ICU-NS groups relative to control neutrophils (n=22, 13, and 26 for ontrol, ICU-S, ICU-NS; respectively). (F) Fluorescence imaging was used to assess annexin V staining in freshly olated neutrophils from all groups. Images were obtained using the Cytation 5 Cell Imaging Multi-Mode eader (Agilent) and analysis was performed using Gen5 Software. Scale bar: 100 µm. (G) Cellular distribution f annexin fluorescence in all groups confirms significant decrease in apoptotic neutrophils (n=7, 6, and 11 for ontrol, ICU-S, and ICU-NS; respectively).

### Differential expression of miRNAs in isolated neutrophils points at impaired clearance of non-survivors’ neutrophils

In an attempt to unravel causes of neutrophilia and to have deeper insights on altered molecular factors that regulate metabolic networks in critically ill patients we analyzed the miRNA pattern in purified neutrophils from a small subset of samples (2 ICU-S and 4 ICU-NS, Fig. 2). A total of 31 miRNAs were differentially expressed between the ICU-S and the ICU-NS groups. Twelve miRNAs are significantly upregulated and 19 miRNAs are significantly downregulated (p<0.05) in the ICU-NS relative to the ICU-S, as shown in Fig. 2A. Hierarchical clustering analysis of differentially expressed miRNAs (DEMs) showed that miRNAs with a similar expression pattern were clustered based on mortality outcome. As shown in Fig. 2B, there are significant differences in the miRNA expression between non-survivors and survivors of COVID-19 patients. Enrichment analysis outlines the most important pathways linked to the reported DEMs based on target genes. Pathway significance enrichment analysis revealed that the differentially expressed miRNA target genes have a close relationship with signaling pathways including the Wnt signaling pathway, PI3K-Akt pathway, Jak-STAT pathway, TGF-β signaling pathway and apoptosis pathway. Interestingly, an increase in miR-15a, miR-15b, miR-223-3p, miR-30d-5p, miR-30e-5p, and miR-363-3p was shown to down regulate genes involved in neutrophil apoptosis as shown in Fig. 2C. No transcriptomic evidence substantiating glycolytic reprogramming was detected in isolated neutrophils from non-survivors. Taken together, the differential miRNA profile suggests that impaired apoptotic clearance of neutrophils leading to neutrophilia is predominant especially in non-survivors’ neutrophils.

**Figure 2.**
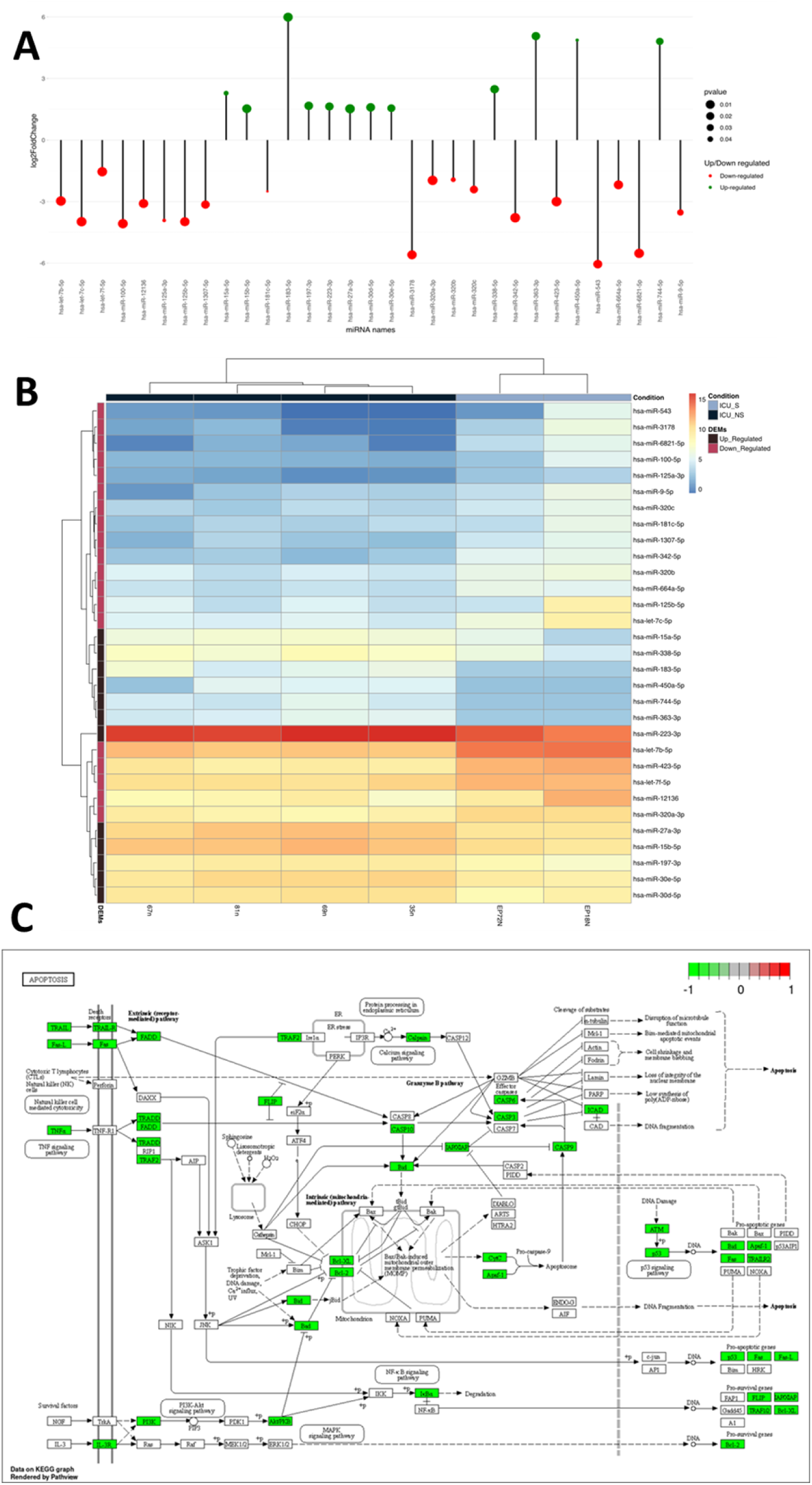
Differential miRNA profile of isolated neutrophils reveals impaired apoptotic clearance of nonsurvivors’ neutrophils. (A) A lollipop plot represents differentially expressed miRNA of non-survivors (n=4) relative to survivors (n=2) purified neutrophils. The x-axis represents miRNA ID, and the y-axis represents log2 fold changes. Each lollipop represents a differentially expressed miRNA between the two groups, with green lollipops indicating upregulated miRNAs and red lollipops indicating downregulated miRNAs. The size of the dot represents the significance level of the change in gene expression. (B) Heat map of 31 differentially expressed miRNAs between survivors and non-survivors COVID-19 patients. Each column represents an individual patient and rows listing different miRNAs that are color-coded to reflect miRNA abundance in each sample. Red indicates a high miRNA amount in the sample, and blue indicates a low gene expression in the sample. The clustering on the top and left of the heatmap show miRNAs similarity in expression patterns. The annotations for each sample indicate the group to which it belongs ICU-S indicating survivors and ICU-NS indicating non-survivors. (C) Apoptosis pathway visualization illustrate the down regulated effect of explored miRNA with its target genes.

### Lung damage and neutrophil count predict mortality in critically ill patients

SARS-CoV-2 infection usually leads to complications such as pneumonia, and in severe cases, ARDS and sepsis, which are now known to leave lasting damage to the lungs including interstitial lung disease [38]. Since activated neutrophil infiltration into inflamed lung is a hallmark of ARDS [5], we analysed the extent of lung damage in a subset of the studied cohort (Fig. 3). We analyzed retrieved chest radiographs for 23 ICU-S and 45 ICU-NS as described in Methods; representative images are given in Fig. 3A. ICU-NS group showed a significantly greater lung damage with a mean score ± SD = 15.93 ± 5.87 vs. 11.61 ± 4.68 for ICU-S group (p=0.003), Fig. 3B. We then proceeded to explore the prognostic power of the CT score as a mortality predictor. We started by generating a receiver-operating characteristic (ROC) curve for the prediction of mortality in terms of the CT score (Fig. 3C). This analysis established a CT damage score cut-off value of 13.5 (AUC = 0.745, p=4×10^-4^) as a stratifying parameter for mortality prediction in the subsequent Kaplan-Meier survival analysis (Fig. 3D). CT lung damage score ≥ 13.5 was subsequently used as a predictor of mortality within 20 days of image acquization (31 out of 38, 81.5%; Log Rank χ2 = 7.26, p=0.007). We have shown recently that neutrophil count is a good severity predictor during the course of COVID-19 infection [16]. Here we tested if neutrophilia can be a mortality predictor for the studied cohort of severely ill patients. The ROC curve for neutrophil counts (% total) established a cut-off value of 69.78 % (Fig. 3E, AUC = 0.798, p=9.2×10^-9^) which was then used in Kaplan-Meier survival analysis in Fig. 3F. Group with neutrophil counts greater than 69.78% of blood lysate showed higher mortality (40 out of 53, 75.5%; Log Rank χ2 = 29.2, p=6.5×10^-8^), Fig. 3F.

**Figure 3.**
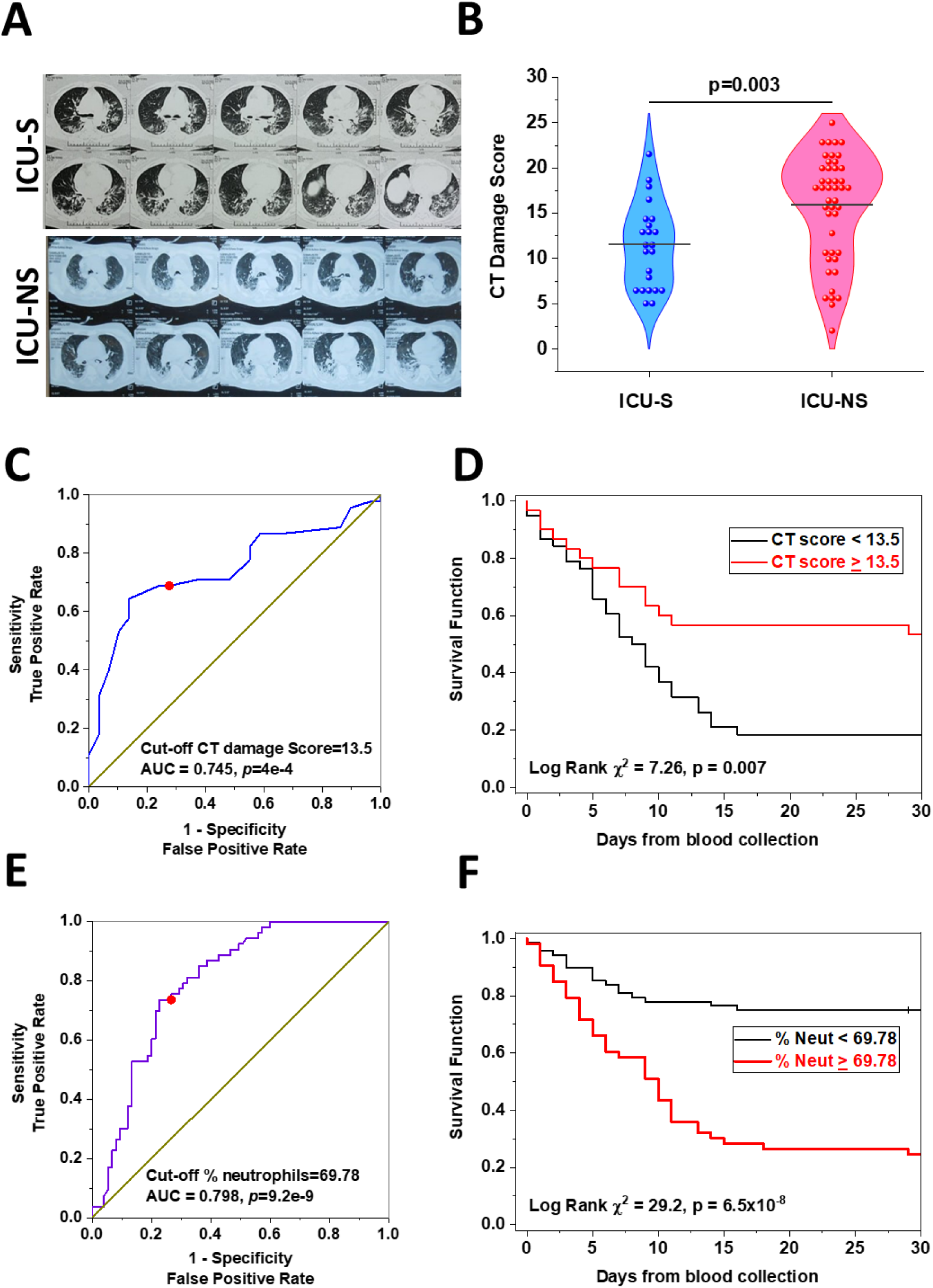
Association of lung damage and neutrophil count with mortality outcome in ICU-COVID-19 ubjects. (A) Representative chest radiographs for ICU-S and ICU-NS groups. (B) A violin plot showing a gnificant increase in lung damage in the ICU-NS group relative to the ICU-S group (n=23 and 45 for ICU-S and U-NS; respectively). (C, E) Receiver operating characteristics (ROC) showing optimal sensitivity and specificity of (C) CT lung damage score, and (E) neutrophil counts as predictors of mortality in severe COVID-19 patients, ith cut-off values and statistical results given. (D, F) Kaplan–Meier estimates of time-to-mortality from blood ample collection during ICU hospitalization. Log-rank Kaplan–Meier survival analyses were carried out to estimate the probability of survival of critically ill patients in relation to cut-off thresholds determined through he ROC analysis.

### Non-survivors’ hemoglobin exhibit reduced oxygen carrying function

So far, our results suggest that both neutrophil count and lung damage are reasonably good mortality predictors in the context of severe COVID-19 illness. The PaO2/FiO2 ratio is frequently used to determine the severity of lung injury in mechanically ventilated patients. Here we show that the PaO2/FiO2 ratio is remarkably lower in the case of ICU-Non-survivors, indicating a marked hypoxemia (Fig. 4A, ICU-S: 319.9 ± 155.9; n=13, ICU-NS: 122.0 ± 110.7; n=26. ICU-S vs. ICU-NS: p= 5.1×10^-5^). Oxygen-hemoglobin dissociation curves (ODC) were constructed using ICU-Survivors (ICU-S, n=26) and ICU-Non-survivors (ICU-NS, n=57) COVID-19 patient data and compared to the standard human ODC at T = 37°C, pH = 7.4, created by the Severinghaus model (Fig. 4B). First, the theoretical standard human blood ODC was estimated based on the original computations implemented by Severinghaus [39]. A left shift of the Hemoglobin oxygen dissociation curve of the ICU-S group is observed, while Hemoglobin oxygen dissociation curve of the ICU-NS group is shifted towards higher oxygen pressure to reach the same degree of hemoglobin saturation. These results suggest an exacerbated respiratory dysfunction and relatively impaired gas exchange in ICU-Non-survivors COVID-19 patients.

**Figure 4.**
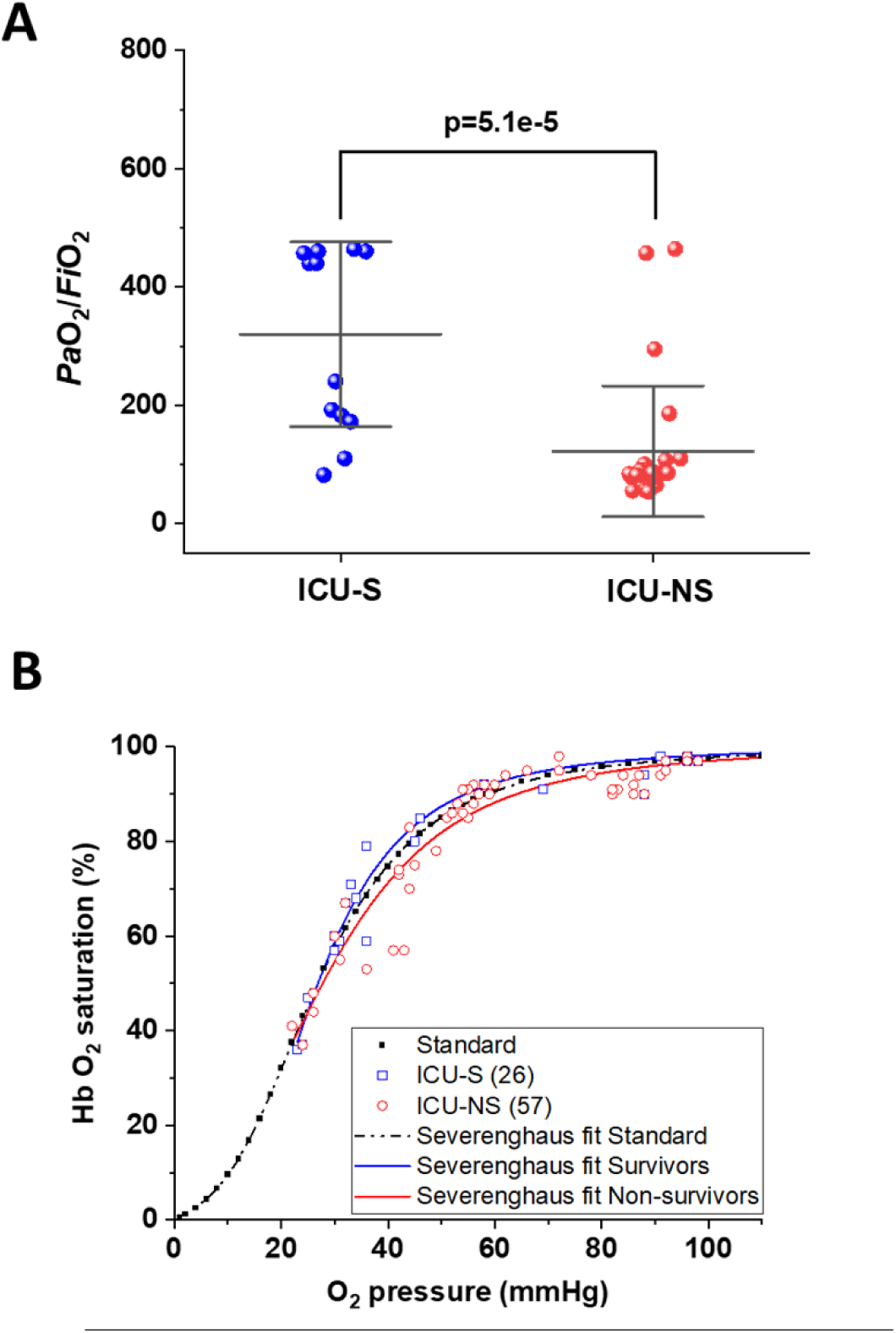
Impaired oxygen carrying function of hemoglobin in non-survivors. (A) A diagram showing a significant decrease in PaO2/FiO2 ratio in ICU-NS group compared to ICU-S group ((n=13 and 26 for ICU-S, and ICU-NS; respectively)). (B) Graphic depiction of Hb-O2 affinity ODC, demonstrating the standard Severinghaus curve (dashed black curve). Values of oxygenation saturation plotted versus oxygen pressure for ICU-S COVID-19 patients (blue rectangles) and ICU-NS COVID-19 patients (red circles).

### Non-survivors’ blood is more acidic and their plasma exhibit hyperlactatemia

Next, we reasoned that the observed shift in the ODC may be due to lowered blood pH during the course of the critical COVID-19 illness especially in non-survivors. Analysis of the chemistry laboratory results retrieved during hospitalization revealed that the average blood pH value was indeed lower in non-survivors (pH = 7.37 ± 0.12, n=53) relative to survivor group (7.43 ± 0.06, n=29, p=0.01) (Fig. 5A). Declines in blood pH can be a result of either a respiratory or a metabolic acidosis. Respiratory acidosis is caused by hypoventilation, and subsequent elevation of PCO_2_. Metabolic acidosis is the result of excessive generation of acid, most commonly lactic acid and/or enhanced loss of HCO_3_^−^ [40]. Laboratory results obtained for participants at the site of sample collections during hospitalization revealed non-significant differences in PCO_2_ (ICU-S, n=29; ICU-NS, n=53), [HCO_3_^−^] (ICU-S, n=30; ICU-NS, n=54), or in hemoglobin levels (ICU-S, n=39; ICU-NS, n=60) between ICU-Survivors and ICU-Non-survivors (supplementary Fig. S1). However, plasma levels of lactate determined colorimetrically, were significantly higher in ICU-NS patients than in controls (Control: 5.62 ± 1.5; n=14, ICU-S: 6.34 ± 2.32; n=13, ICU-NS: 7.47 ± 2.2; n=21. ICU-S vs. Control: p=0.064; ICU-NS vs. Control: p= 0.032; ICU-S vs. ICU-NS: p=0.27), indicating a marked lactic acidosis in the non-survivors’ group (Fig. 5B). To further confirm these findings, we also analyzed blood levels of lactate in stored blood samples from a subset of controls, ICU-Survivors and ICU-Non-survivors subjects using a blood lactate meter (Control: 3.55 ± 1.4; n=11, ICU-S: 5.51 ± 2.01; n=15, ICU-NS: 7.37 ± 3.19; n=15. ICU-S vs. Control: p=0.11; ICU-NS vs. Control: p= 7.7×10^-4^; ICU-S vs. ICU-NS: p=0.1). Our results showed that blood lactate levels were significantly elevated in the ICU-NS group when compared with controls (Fig. 5C).

**Figure 5.**
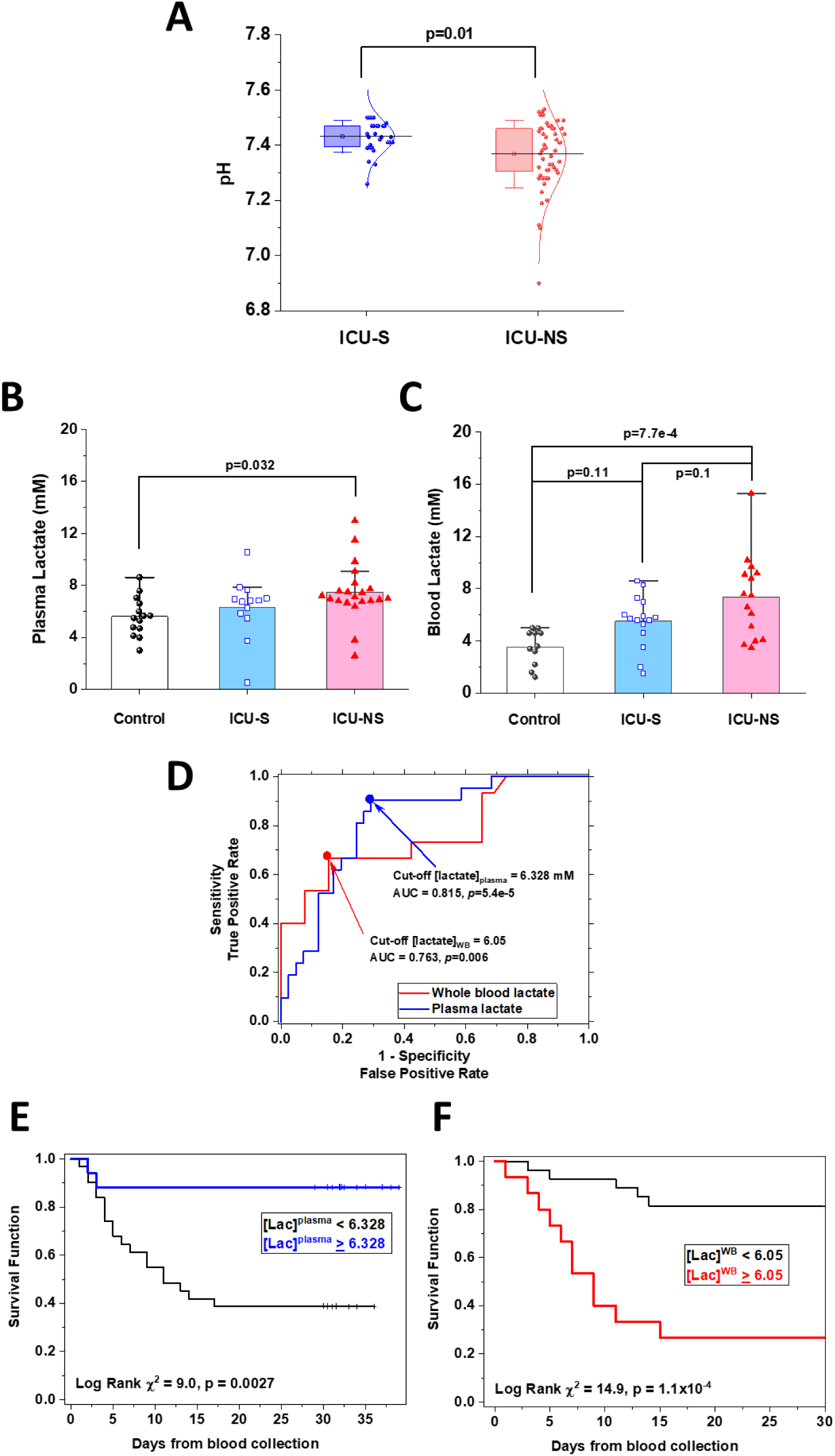
Blood acidosis and hyperlactemia in non-survivors and association of plasma and blood lactate levels with mortality outcome in ICU-COVID-19 subjects. (A) A box plot showing a significant decrease in blood pH in ICU-NS group when compared to the ICU-S group (n=29 and 53 for ICU-S, ICU-NS; respectively). (B) box plot showing significant increase in plasma lactate level in ICU-NS group when compared to control group (n=14 and 21 for control, ICU-NS; respectively). (C) A box plot showing significant increase in blood lactate level in ICU-NS group when compared to the control group (n=11 and 15 for control, ICU-NS; espectively). (D) Receiver operating characteristics (ROC) showing optimal sensitivity and specificity of plasma nd blood lactate levels as predictors of mortality in severe COVID-19 patients, with cut off values [6.328 (AUC = 0.815, p=5.4 x 10-5) and 6.05 mM (AUC = 0.763, p=0.005) for plasma and blood lactate levels, respectively]. E-F) Kaplan–Meier estimates of time-to-mortality from blood sample collection during ICU hospitalization were stratified per the cut-off values obtained through the ROC analyses.

### Lactate levels in blood and plasma predict mortality in critically ill patients

Next, we explored if whole blood and plasma lactate could predict mortality outcome in severe patients (Fig. 5D-F). An analysis of Kaplan–Meier estimates of time-to-mortality following blood sample collections during ICU hospitalization in relation to plasma and blood lactate was performed. Receiver operating characteristic curves (ROC) were used to determine the optimal sensitivity and specificity of plasma and blood lactate (Fig. 5D). ROC-determined cut-off values for each of these parameters were: plasma lactate, 6.328 mM (AUC = 0.815, p=5.4 x 10^-5^); and blood lactate concentration, 6.05 mM (AUC = 0.763, p=0.006). Finally, we stratified patients into two groups: patients showing values below or equal to the cut-off values versus those exhibiting values above the cut-off values for each of the studied parameters. Kaplan-Meier analysis indicated that patients with plasma lactate ≥ 6.328 mM showed significantly higher in-hospital mortality (19/31, 61.3%, logrank χ2 = 9.0, p = 0.0027, Fig. 5E). Similarly, patients with higher lactate in their blood showed higher mortality (11/15, 73.3%, logrank χ2 = 14.9, p = 1.1 x 10^-4^, Fig. 5F).

### COVID-19 severity and mortality are not associated with significant glycolytic shift in individual neutrophils, but neutrophilia significantly contributes to lactate production in non-survivors

We asked if and how leukocytes contribute to the observed increase in blood/plasma lactate levels and lowered pH, especially in non-survivors (Fig. 6). Lactate was found to be released by human neutrophils as a major byproduct of glycolysis [17] and increasing neutrophil counts is a hallmark of COVID-19 severity and mortality (Figs. 1B and 3F). To identify potential contributors to lactic acidosis in circulating blood, we assessed and compared glycolytic activities in isolated neutrophils, platelets, and peripheral blood mononuclear cells (PBMCs) from the current cohort of critically ill patients relative to the control group. First, group-dependent changes in cell numbers in these three cell types were assessed using flow cytometry to show that only neutrophil counts exhibit a remarkable increase with COVID-19 infection (Fig. 6A, Control: 499.32 ± 338.9; n=9, ICU-S: 1121.22 ± 952.3; n=18, ICU-NS: 1491.76 ± 1168.74; n=25. ICU-S vs. Control: p=0.29; ICU-NS vs. Control: p= 0.04; ICU-S vs. ICU-NS: p=0.46). In fact, PBMCs showed a significant decrease in non-survivors, Fig. 6A (Control: 299.43 ± 132.25; n=7, ICU-S: 175.71 ± 146.07; n=8, ICU-NS: 117.34 ± 122.75; n=10. ICU-S vs. Control: p=0.19; ICU-NS vs. Control: p= 0.03; ICU-S vs. ICU-NS: p=0.63).

**Figure 6.**
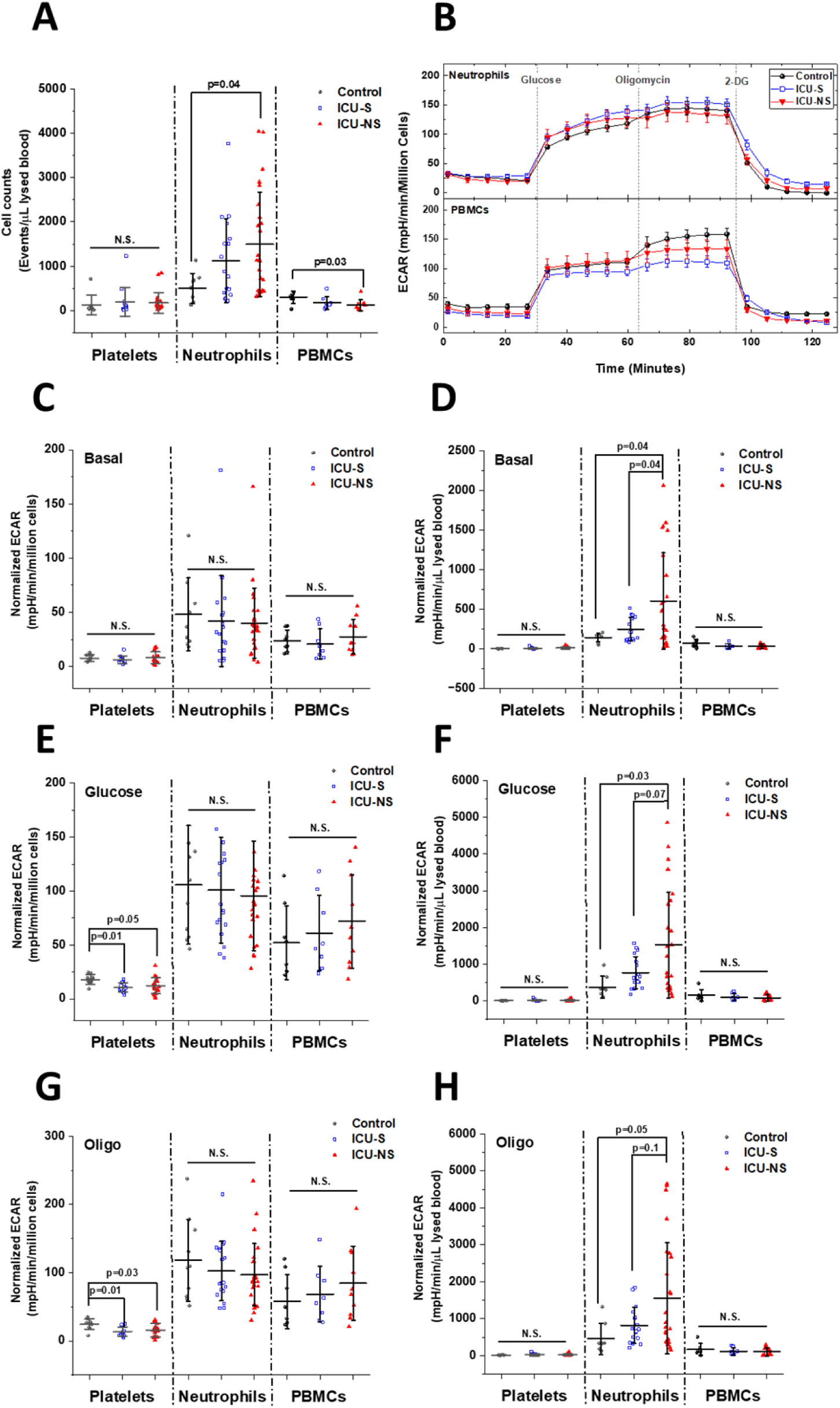
A significant increase in the neutrophil-mediated glycolytic lactate production is only seen when neutrophilia is considered. (A) Only non-survivors’ neutrophils show a significant increase in the overall count er μL of lysed blood as detected by flow cytometry. Both platelet and PBMCs counts show no dependence on mortality. (B) Representative seahorse traces for ECAR (glycolytic flux) measured in neutrophils and PBMCs freshly isolated from control, ICU-S, or ICU-NS group and normalized to seeded count. (C, E, G) When basal CAR, glucose-supplemented glycolytic flux, and maximal glycolytic capacity were normalized to seeded cell counts, none of the studied cell types showed dependence on the mortality state except for platelets, which showed a consistent decrease in the ICU-S and ICU-NS groups. (D, F, H) When these ECAR values were multiplied by cell counts in the μl-lysed blood, only ICU-NS neutrophils displayed a significant increase relative o ICU-S and control groups. The number of subjects analyzed were as follows: Platelets, (n=8, 14 and 20 for control, ICU-S and ICU-NS groups; respectively); neutrophils, (n=10, 14, and 20 for control, ICU-S, and ICU-NS groups; respectively); and PBMCs, (n=8, 8 and 10 for control, ICU-S and ICU-NS groups; respectively).

Traces of Seahorse metabolic analysis of freshly isolated neutrophils from subsets representing all groups are shown to demonstrate the independence of the extracellular acidification rate by individual neutrophils on mortality outcome (Fig. 6B). Generally, for both neutrophils and PBMCs, basal ECAR, glycolytic metabolism, and glycolytic capacity normalized to seeded cell counts did not show any significant differences among all groups (Fig. 6, panels C, E, G). Only platelets exhibited significantly reduced glycolytic flux in critically ill COVID-19 patients, which confirms our recently published data [22]. However, a significant increase in global basal ECAR per μL lysed blood (Fig. 6D, Control: 144.07 ± 53.94; n=8, ICU-S: 248.91 ± 148.41; n=17, ICU-NS: 604.55 ± 609.41; n=25. ICU-S vs. Control: p=0.85; ICU-NS vs. Control: p= 0.04; ICU-S vs. ICU-NS: p=0.04), glucose-supplemented glycolytic metabolism (Fig. 6F, Control: 378.13 ± 296.94; n=8, ICU-S: 768.70 ± 436.25; n=17, ICU-NS: 1525.42 ± 1439.79.74; n=25. ICU-S vs. Control: p=0.67; ICU-NS vs. Control: p= 0.03; ICU-S vs. ICU-NS: p=0.07), and glycolytic capacity when mitochondria are inhibited with oligomycin (Fig. 6H, Control: 455.51 ± 423.35; n=8, ICU-S: 819.45 ± 491.12; n=17, ICU-NS: 1559.55 ± 1505.89; n=25. ICU-S vs. Control: p=0.73; ICU-NS vs. Control: p= 0.05; ICU-S vs. ICU-NS: p=0.10) have been detected in ICU-NS neutrophils relative to ICU-S and control groups. These results indicate that the increase in glycolysis in ICU-NS neutrophils is due to a fundamental increase in neutrophil counts rather than being caused by a metabolic shift in individual neutrophils.

### Neutrophil counts associate with blood pH and plasma lactate levels

We then investigated the relationship between neutrophil counts and lactic acidosis in COVID-19 ICU-patients. A rise in lactate levels that coincides with increased neutrophils counts in the circulation and inflamed tissues has been demonstrated in multiple pathological conditions including shock, sepsis, and ischemia [41] but not in COVID-19. Here, we compared neutrophil counts with blood pH in critically ill patients as shown in the contoured scatter plot (Fig. 7A) which shows an increased tendency towards lower pH in patients exhibiting greater neutrophil counts. In Fig. 7B we show that blood lactate levels positively correlate with neutrophil counts (Pearson’s r=0.43, p=0.009), a trend that was also conserved for plasma lactate levels albeit with weaker but significant correlation (Fig. 7C, Pearson’s r=0.36, p=0.017). These results further suggest that neutrophilia are significantly contributing to lactic acidosis in critically ill patients. These results recalled an important question: Is there a link between neutrophilia and lung damage? To answer this question, we plotted the CT damage score as a function of neutrophil counts (Fig. 7D). Although no statistically significant correlation has been observed to linearly connect these parameters, a qualitative association can be seen that indicates a denser contour distribution at higher values of both neutrophil counts and CT damage scores. This suggests a subtle association between neutrophilia and COVID-19-caused lung damage.

**Figure 7.**
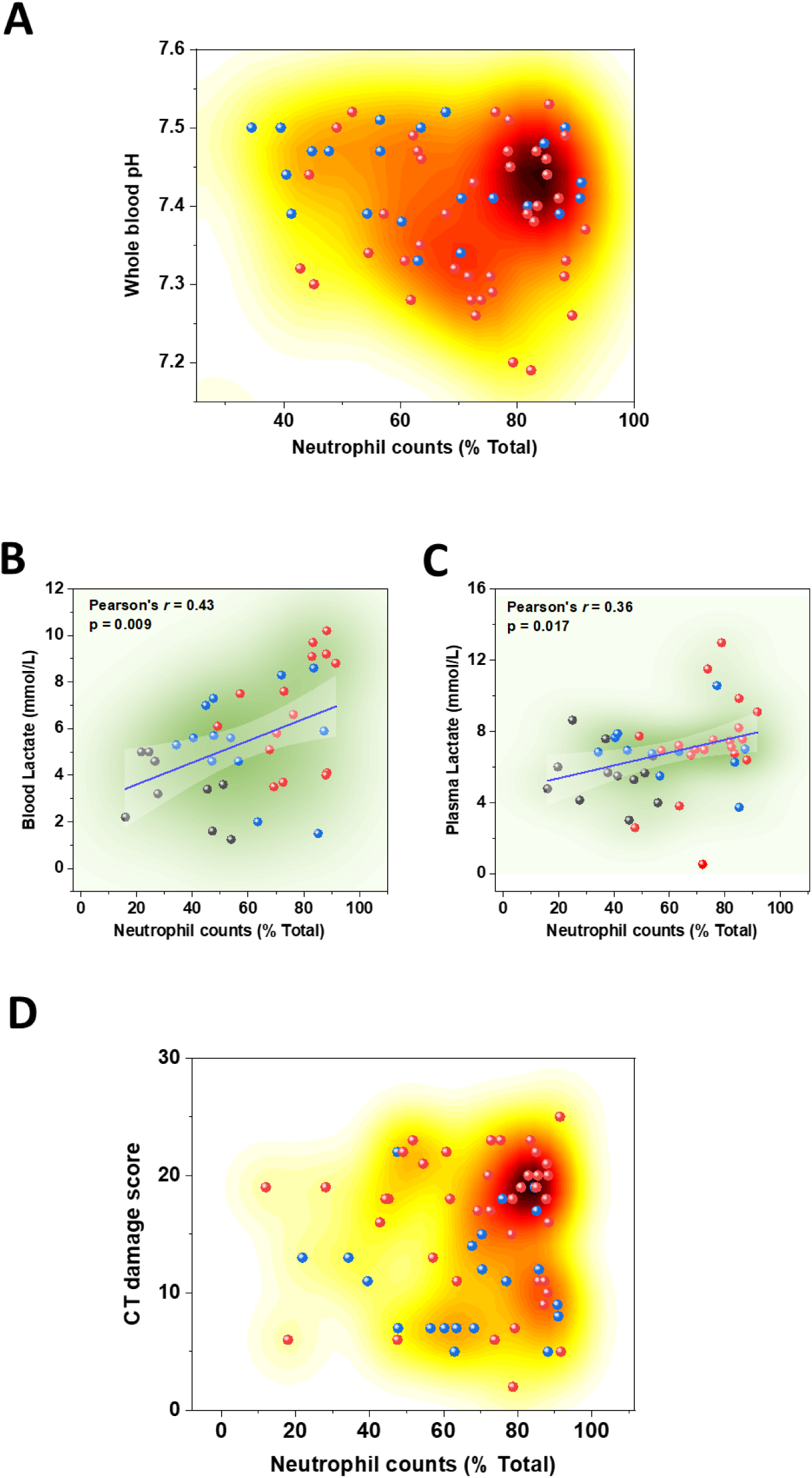
Associations of neutrophilia with blood acidosis and hyperlactatemia in blood and plasma of critically ill subjects. (A) Colored-contour map reflecting the density distribution of whole blood pH values in relation to neutrophil counts in critically ill COVID-19 patients. The density map suggests that patients with higher neutrophil counts tend to possess more acidic blood. (B, C) Linear regressions showing that both blood and plasma lactate increase with neutrophil counts. Results of linear least square fitting along with confidence intervals are superposed on density maps to show data distributions. (D) Colored-contour map reflecting the density distribution of CT lung damage scores in relation to neutrophil counts of critically ill COVID-19 patients. Tendency of increased lung damage at higher neutrophil counts are readily observable. Availability of paired data from individual patients dictated the number of points used in each correlation.

## Discussion

Our COVID-19-related research efforts have been focused on a specific question: in a relatively homogeneous cohort of intensive care hospitalized severe patients that exhibit somewhat similar demographic and clinical characteristics, why does a differential mortality outcome arise? We implicated neutrophils in extensive oxidative damage of circulating human serum albumin [16] and in platelets activation and hypercoagulability [22] especially in patients with poor mortality outcome. Neutrophilia in critical COVID-19 illness and mortality outcome is among the earliest risk factors identified [42]. However, substantial research efforts are still required to unravel the regulatory mechanisms of neutrophilia in COVID-19. Here we highlight the contribution of neutrophilia to a multifaceted array of COVID-19-associated cellular and systemic disorders including lung damage, metabolic remodeling, acidosis, and oxygen delivery during the climax of infection.

The production and clearance of neutrophils are tightly regulated to maintain homeostasis, both in physiologic as well as pathologic conditions. In healthy individuals, neutrophil production is primarily controlled by the cytokine granulocyte-colony stimulating factor (G-CSF), which stimulates the bone marrow to produce and release neutrophils into the circulation. Neutrophils have a short lifespan and are cleared from the circulation by apoptosis and subsequent phagocytosis by macrophages and other phagocytic cells. In pathological conditions such as infection, trauma, or inflammation, neutrophil production and activation are increased to combat the underlying condition. However, excessive or dysregulated neutrophil activation can lead to tissue damage and contribute to the pathogenesis of various diseases, including autoimmune disorders and chronic inflammatory conditions. Therefore, maintaining proper neutrophil homeostasis is critical for maintaining a healthy immune system and preventing the development of disease [reviewed in [43].

This work adds to the bulk of literature reporting increased neutrophil counts in the context of severe COVID-19 infection. Furthermore, we provide new evidence from transcriptomic analysis in purified neutrophils that genes underlying apoptotic machinery are significantly down-regulated in non-survivors albeit with a mechanism that awaits further exploration. Concomitantly, we showed that circulating neutrophils are remarkably less apoptotic in critically ill patients especially non-survivors. Consequently, impaired apoptosis seems to contribute to the accumulation of neutrophils leading potentially to the exacerbation of neutrophil histotoxic destructive capacity of tissues and organs [43]. Not surprisingly, we found that mortality outcome increased in patients exhibiting more extensive lung damage and increased neutrophil counts. This suggests that targeting neutrophil apoptosis in the context of COVID-19 infection might be a viable therapeutic pathway to combat excessive innate immune activation and organ damage in severe cases.

In this study, we also show that the oxygen dissociation curve of the ICU-NS group exhibits a right shift when compared with that at standard condition or of the ICU-S groups. Published data on ODC curves stratified per mortality outcome in hospitalized critically ill COVID-19 patients are scarce (reviewed recently in [44]) but the few identified studies suggest that left-shifted ODCs are usually associated with better prognosis [45]. Recently, it has been proposed that ODC in critically ill COVID-19 patients display a left shift when compared with the standard ODC [46], or with patients suffering from hypoxia caused by other respiratory disorders [47]. Indeed, our results indicated that ODC of ICU-Survivors COVID-19 patients is shifted to the left compared with the standard condition, while the ODC of ICU-Non-survivors is shifted towards higher oxygen pressure to reach the same degree of hemoglobin saturation. These results are in concordance with the recent study by Ceruti et al., which showed an absence of left shift in critically ill COVID-19 patients with poor outcomes, compared with those discharged from the ICU [46].

Decreasing oxygen affinity, which hinders O_2_ uptake in the lungs is detrimental and was speculated to result from elevated core body temperature in patients with poor prognosis [48]. Here we tested an alternative hypothesis implicating lowered pH as a cause of lowered oxygen affinity by hemoglobin in patients with poor mortality outcome. Numerous studies have presented evidence that COVID-19 severity and high in-hospital mortality are often associated with severe hypoxemia, hyperlactatemia and acidosis. In fact, oxygen deprivation of tissue provokes the transition to anaerobic metabolism and enhances the production of lactate [9]. Elevated lactate feeds back into this cycle by lowering the average pH and reducing the oxygen carrying capacity of hemoglobin [49]. However, the sources underlying the overproduction of lactate leading to lactic acidosis in critically ill COVID-19 patients in relation to mortality outcome are still unexplored experimentally. Dramatic increases in neutrophil counts in ICU-hospitalized patients enticed us to explore the subsequent implications of their metabolic remodeling while hypothesizing a role for lactate overproduction by glycolytic neutrophils in the well-documented impairment of oxygen delivery and hypoxemia.

Indeed, our analysis indicated that non-survivors’ blood is generally more acidic, and that this is associated with significantly increased blood and plasma lactate concentrations. Our results are in tune with the study by Kieninger et al., which identified low blood pH as a significant prognostic factor for in-hospital mortality in critically ill COVID-19 patients [50]. The observed drop in blood pH can be a consequence of either respiratory or metabolic acidosis. Respiratory acidosis is caused by hypoventilation, and subsequent elevation of PCO_2_, whereas metabolic acidosis is the result of excessive generation of acid, most commonly lactic acid and/or enhanced loss of HCO_3_^−^ [40]. In this context, we tested if mortality is associated with increased blood PCO_2_ or HCO_3_^−^. Our results showed non-significant differences in PCO_2_ or HCO_3_^−^ levels between ICU-Survivors and ICU-non-survivors. In contrast, ICU-Non-survivors displayed significantly higher levels of plasma and blood lactate compared with ICU-Survivors, pointing to a significant lactic acidosis in critically ill COVID-19 patients with fatal outcomes. Our findings are consistent with other studies showing that non-survivors covid-19 patients exhibited greater levels of blood lactate when compared with survivors (reviewed in: [14]). Hypoxia and acidosis reinforce each other in a viscous cycle. According to the Verigo-Bohr effect, a slight decline in blood pH causes a significant reduction in oxygen saturation. With the progression of oxygen insufficiency, acidosis is exacerbated and thus continues to hinder oxygen delivery to peripheral tissues [9].

We also attempted to shed some light on the source(s) of circulating lactate in the studied patients. Under physiological conditions, red (RBCs) and white (WBCs) blood cells are minor sources of lactate. Although RBCs exclusively rely on glycolysis for their bioenergetic demands, we ruled out RBCs as an important player in the observed metabolic acidosis because RBCs were shown to act as a lactate sink in COVID-19 [51]. Furthermore, we have previously shown that glycolytic activities in platelets actually decrease with severity [22]. However, the situation with the WBCs might be different during immune activation and inflammation, presumably due to over-activation and metabolic rewiring of the activated WBCs. We thus characterized glycolytic activities in platelets, neutrophils, and PBMCs freshly isolated in representative groups of controls, survivors, and non-survivors COVID-19 patients. Surprisingly, we found that COVID-19 severity and mortality are not associated with glycolytic shifts in any of the studied cell types except for a pronounced reduction in glycolytic fluxes in platelets. However, when we considered the changes in cell counts for each blood cell type, we concluded that neutrophilia significantly contributes to lactate production in non-survivors. We also demonstrated that elevated plasma and blood lactate levels as well as blood acidity are significantly correlated with enhanced neutrophil counts in non-survivors COVID-19 patients.

## Conclusion

Collectively, the current findings confirm that neutrophilia are significantly contributing to lactic acidosis, impaired oxygen delivery, and lung damage in critically ill patients. These findings may constitute an additional step towards understanding the pathophysiology of lactic acidosis detected in critically ill COVID-19 patients with fatal outcomes. Our data also suggests that targeting neutrophil apoptosis in the context of COVID-19 infection might be a viable therapeutic pathway to combat excessive innate immune response and organ damage in severe cases.

### List of Abbreviations

ICU-S: Intensive Care Unit-Survivors
ICU-NS: Intensive Care Unit-Non-survivors
CT: Computed Tomography
ROC: Receiver operating characteristic curves
ARDS: Acute Respiratory Distress Syndrome
RT-PCR: Real Time Polymerase Chain Reaction
HBSS: Hank’s Balanced Salt Solution
PBS: Phosphate Buffer Saline
PBMCs: Peripheral Blood Mononuclear cells
ECAR: Extracellular acidification rates
DEMs: Differentially Expressed miRNAs
G-CSF: Granulocyte-colony stimulating factor
ODC: Oxygen Dissociation Curve
RBCs: Red Blood Cells
WBCs: White Blood Cells

## Declarations

### Ethics Approval

The studies involving human participants were reviewed and approved by Children’s Cancer Hospital’s Institutional Review Board. The patients/participants provided their written informed consent to participate in this study.

### Availability of data and materials

The original contributions presented in the study are included in the article/Supplementary Material. Further inquiries can be directed to the corresponding authors.

### Disclosure of Conflicts of Interest

The authors declare that the research was conducted in the absence of any commercial or financial relationships that could be construed as a potential conflict of interest.

### Funding

The present work was funded by the Association of Friends of the National Cancer Institute and the Children’s Cancer Hospital Foundation.

### Authorship Contributions

BAY and AAE performed experiments, analyzed results, and assisted in the manuscript preparation. Hel-S, MZ, AGK, RH, MAB, and AS helped in data collection. Hel-Sh, OS, and AS performed the transcriptomic analysis. MSH provided materials and analyzed clinical data. Subjects’ recruitment, ethical approvals, clinical follow-up and assessments of lung damage were done by RE-M and ME-A. EAA-R conceived and directed the project, helped design experiments and wrote the initial draft of the manuscript, and SSA conceived and planned the whole project, designed all experiments, obtained funding and committee approvals, analyzed data, prepared figures and wrote the final manuscript.

## Acknowledgement

Not Applicable

---

**Supplementary figure S1.**
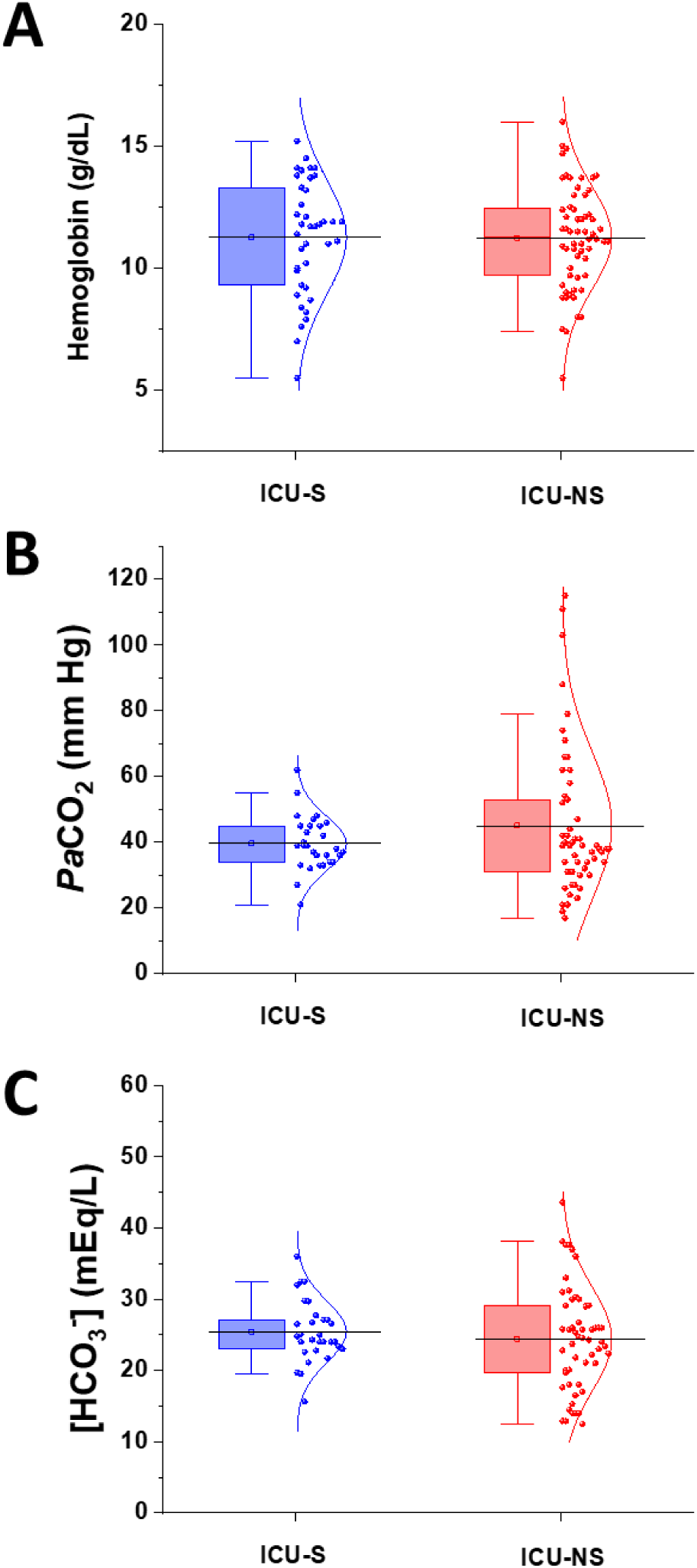
Box plot representations showing non-significant differences in (A) hemoglobin (n=29 and 60), (B) PaCO_2_ (n=29 and 53), (C) [HCO_3_^−^] (n=30 and 54) levels between ICU-S and ICU-NS groups.

## Notes

### Competing Interest Statement

The authors have declared no competing interest.

